# Asexual reproduction drives the reduction of transposable element load

**DOI:** 10.1101/501361

**Authors:** Jens Bast, Kamil S. Jaron, Donovan Schuseil, Denis Roze, Tanja Schwander

## Abstract

Theory predicts that sexual reproduction can both facilitate and restrain transposable element (TE) accumulation by providing TEs with a means of spreading to all individuals in a population and facilitating TE load reduction via purifying selection. By quantifying genomic TE loads over time in experimental sexual and asexual *Saccharomyces cerevisiae* populations, we provide direct evidence that asexual reproduction drives a reduction of TE loads. We show, using simulations, that this reduction occurs via evolution of TE activity, most likely via increased excision rates. Thus, sex is a major driver of genomic TE loads and at the root of the success of TEs.

## Main Text

Self-replicating transposable elements (TEs) can occupy large fractions of genomes in organisms throughout the tree of life (reviewed in Hua-Van et al., 2011). Their overwhelming success is driven by their ability to proliferate independently of the host cell cycle via different self-copying mechanisms (i.e., in a ‘cut-and-paste’ or ‘copy-and-paste’ style). These mechanisms allow TEs to invade genomes similarly to parasites despite generally not providing any advantage to the individual carrying them (Doolittle and Sapienza, 1980; Orgel and Crick, 1980). To the contrary, TEs generate deleterious effects in their hosts by promoting ectopic recombination and because most new TE insertions in coding or regulatory sequences disrupt gene functions (Finnegan, 1992; Montgomery et al., 1991).

Theory predicts that sexual reproduction can both facilitate and restrain the genomic accumulation of TEs and it is currently unclear whether the expected net effect of sex on TE loads is positive or negative. Sexual reproduction can facilitate the accumulation of TEs because it allows TEs to colonize new genomes and spread throughout populations (Hickey, 1982; Zeyl et al., 1996). Because the colonization of new genomes is more likely for active TEs, sexual reproduction should favor the evolution of highly active TEs (Charlesworth and Langley, 1986; Hickey, 1982), even though increased activity generates higher TE loads and additional deleterious effects in the host genome. At the same time, sexual reproduction can restrain TE accumulation because it facilitates the evolution of host defences and increases purifying selection against deleterious TE copies (Agren and Wright, 2011; Arkhipova and Meselson, 2005; Crespi and Schwander, 2012; Nuzhdin and Petrov, 2003; Wright and Finnegan, 2001). In the absence of sex, reduced purifying selection can thus result in the accumulation of TEs, unless TE copies get eliminated via excision at sufficiently high rates (Dolgin and Charlesworth, 2006).

To quantify whether the net effect of sexual reproduction on genomic TE loads is positive or negative, we study the evolution of genomic TE loads in experimental yeast (*Saccharomyces cerevisiae*) populations generated in a previous study (McDonald et al., 2016). In McDonald et al., four sexual and four asexual strains originating from the same ancestor strain (W303) were maintained under constant conditions. For sexual strains, a mating event was induced every 90 generations. Sequencing of each strain was conducted at generation 0 and every 90 generations prior to mating (for details see **Methods**, and McDonald et al., 2016). In the present study, we use the published Illumina data to quantify TE loads in each strain for each sequenced generation.

TEs in *S. cerevisiae* are well characterized (Carr et al., 2012; Castanera et al., 2016; Voytas and Boeke, 1992). *S. cerevisiae* TEs consist solely of ‘copy-and-paste’ elements that are flanked by long terminal repeats (LTRs) and are grouped into the families *Ty1-Ty5* (Voytas and Boeke, 1992). The 12.2 Mb genome of the studied yeast strain comprises approximately 50 full-length, active*Ty* element copies, and 430 inactive ones (Carr et al., 2012). Inactive copies comprise truncated elements as well as remnants from TE excisions, which consist of a single LTR (Carr et al., 2012). Excisions are driven by intra-chromosomal recombination between the two flanking LTRs of a TE.

Using different computational approaches to quantify genomic TE loads in experimental yeast strains, we show that sex is required for the success of TEs, as TE loads decrease over time under asexual reproduction. For the first approach, we quantified total TE loads, without distinguishing between active and inactive TEs. This was done by computing the fraction of reads that mapped to a curated *S. cerevisiae* TE library (see **Methods**) for each yeast strain and sequenced generation. This analysis revealed that the total TE load in sexual strains remained constant over 1000 generations, but decreased in asexual strains over time (resulting in a total reduction of 23.5% after 1000 generations; generation effect *P* < 0.001, reproductive mode effect *P* = 0.081, and interaction between generation and mode *P* < 0.001; permutation ANOVA, **Fig. S1**). For the second approach, we focused on active (i.e., full-length) TE copy insertions, because only active TEs can increase genomic TE loads over time. Detecting specific TE insertions by aligning short-read data to a reference genome is difficult and associated with a detection bias towards TEs present in the reference genome. With a pipeline that combines different complementary approaches (see **Methods**), the available sequencing data allowed us to detect 24 out of the 50 full-length insertions present in the reference genome. As with the first approach, we found that the number of (detectable) full-length TE copies remained constant in sexual yeast strains, but decreased in asexual strains over time (generation effect *P* = 0.006, reproductive mode effect *P* =0.033, and interaction between generation and mode *P* < 0.001; permutation ANOVA). In asexual strains, the estimated average number of full-length TEs decreased from approximately 50 to 41 over 1000 generations (**Fig. 1**).

**Fig. 1.**
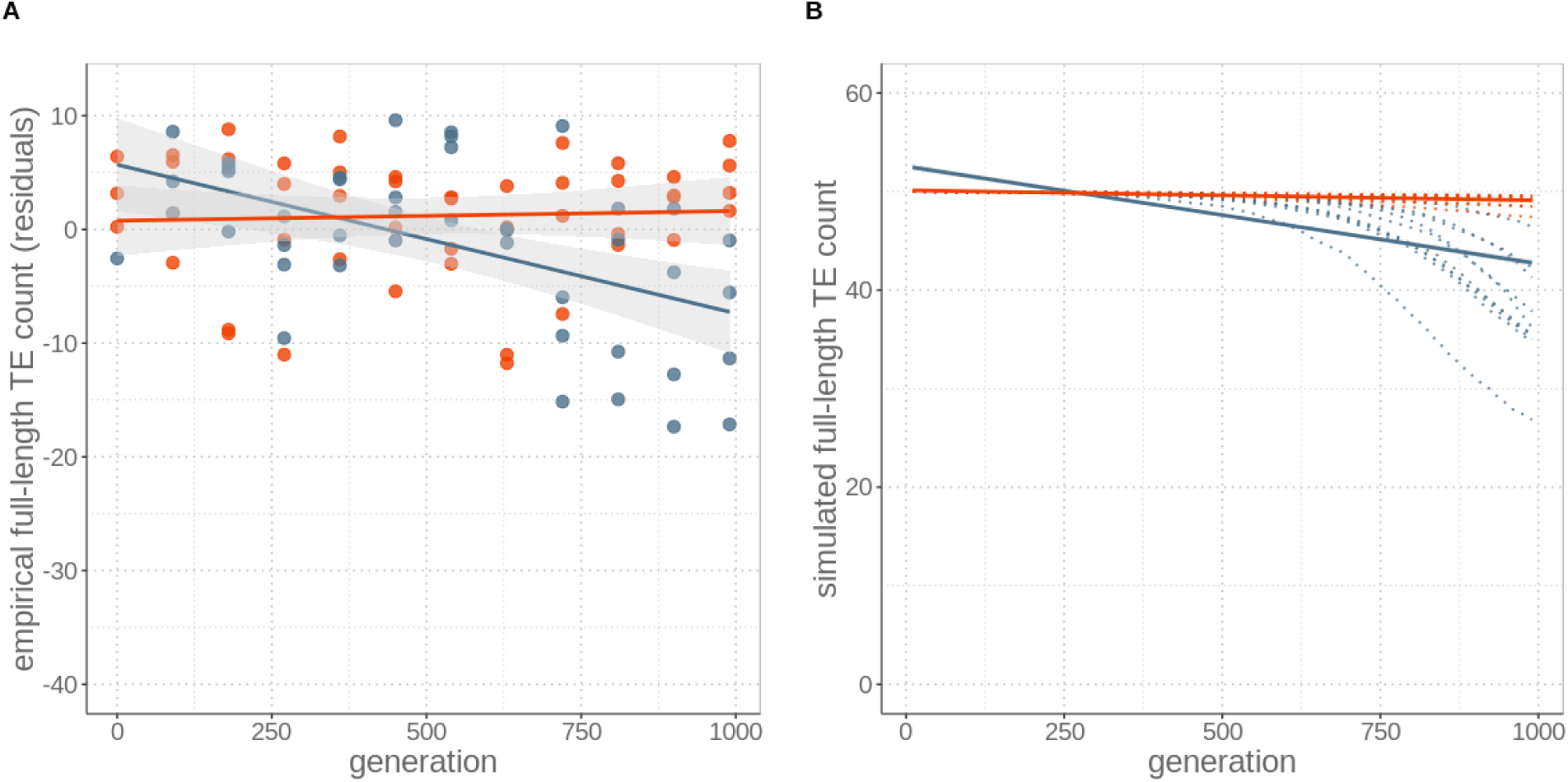
Sex maintains constant TE loads through time, while its absence drives TE copy number reductions, for both (**A**) empirical data and (**B**) simulations including a modifier allele. **(A)** Number of full-length TE copies inserted in genomes for four replicates of sexual (red) and asexual (blue) yeast strains over 1000 generations of experimental evolution. Numbers are expressed as residuals, since the TE detection probability depends on sequencing coverage (Fig. S2). **(B)** Individual-based simulations to study expected TE load dynamics under sexual and asexual reproduction with ten replicates (dotted lines). The simulations are parameterized with yeast-specific values and include an allele modifying TE activity rates. For both (A) empirical and (B) simulation data, asexuals lost about nine active, full-length TEs by generation 1000. Lines represent LR and the grey areas represent 95% CI.

This decrease could be generated by increased TE excision rates in asexual as compared to sexual yeast, reduced transposition rates, or a combination of both mechanisms. To evaluate the relative importance of the two mechanisms, we estimated the number losses of TEs present in the ancestral yeast strain, as well as the number of novel insertions, at each generation (**Fig. 2**). These analyses revealed that ‘ancestral’ TE insertions are lost at a higher rate in asexual than sexual strains (generation effect P = 0.002, reproductive mode effect P = 0.027, and interaction between generation and mode P < 0.001; permutation ANOVA), while we detected similar numbers of novel TE insertions (indicative of similar transposition rates) for the two reproductive modes (generation effect P = 0.338, reproductive mode effect P = 0.271, and interaction between generation and mode P = 0.599; permutation ANOVA). Taken together, our empirical observations indicate that even very rare events of sex (here just 10 out of 990 events of reproduction) are sufficient to maintain genomic TE loads, while asexuality results in the reduction of TE loads, most likely via the evolution of increased TE excision rates.

**Fig. 2.**
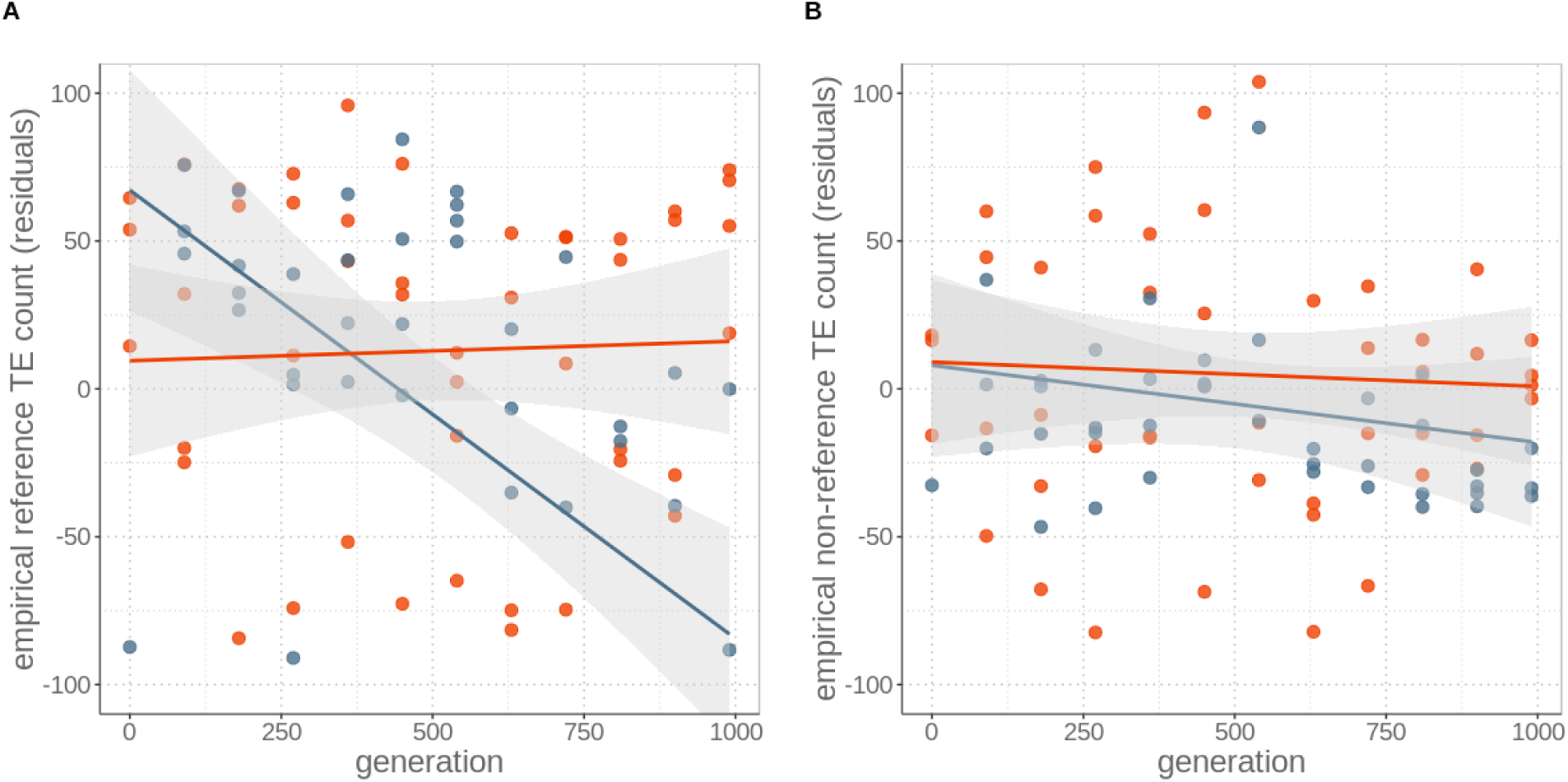
Decrease of insertions in asexuals over time is largely driven by loss of ‘ancestral’ reference insertions (**A**) rather than novel insertions (**B**). Count of any TE insertions, irrespective of full-length TE, solo LTR, truncated elements or other types. Numbers are expressed as residuals, since TE detection probability depends on sequencing coverage. Lines represent LR and the grey areas represent 95% CI.

Our empirical findings suggest that the evolution of reduced TE activity in asexual strains influences genomic TE loads more strongly than purifying selection, which should act to reduce TE loads most effectively in sexual strains. To evaluate the robustness of these findings, we tested whether the net loss of TEs under asexualitly could be recapitualted via a theoretical approach. As explained above, different theoretical approaches have shown that both purifying selection and activity rate evolution can affect TE loads under sexual or asexual reproduction (Charlesworth and Langley, 1986; Dolgin and Charlesworth, 2006; Hickey, 1982). However, there is currently no theoretical framework that studies TE load evolution under the joint effects of the different processes. To fill this gap, we extended the individual-level simulation program of Dolgin and Charlesworth (Dolgin and Charlesworth, 2006). This program allows to study the evolution of TE copy numbers in an asexual lineage as a function of TE activity (the joint effects of transposition and excision rates), as well as of the strength of selection against TE insertions, which depends on the fitness cost per TE insertion. To compare TE loads in sexual and asexual lineages, we first extended the program to include events of sexual reproduction and parameterized the simulations with empirically determined values from yeast (Blanc and Adams, 2004; Carr et al., 2012; Garfinkel et al., 2005; see **Supplementary Material**). We thus ran individual-based simulations with a range of transposition rates, excision rates and selection coefficients with and without epistasis between TE copies as pertinent for yeast (see **Table S2**). For all simulations, TE loads in populations undergoing sex every 90 generations decreased faster than in asexual populations, contrary to our empirical observations. This faster decrease of TE loads in sexual populations occurred because sexual events generated variation among individuals in TE loads (i.e., fitness), which facilitates selection against deleterious TEs (see also Dolgin and Charlesworth, 2006). Implementing different transposition rates under meiosis (sex) or mitosis (asex) did not affect this finding. Indeed, increased TE activity during meiosis only transiently increases TE loads in sexual strains. Because such activity also generates increased variation in TE loads (and therefore fitness) among strains, the additional TE copies generated during meiosis are rapidly removed by purifying selection (**Fig. S4**). In a second step, we therefore allowed for the evolution TE activity rates over time. This was implemented by introducing a modifier allele that increases excision rates. The allele itself has no direct fitness effect, such that it can only be fixed in a population via genetic hitchhiking. In simulations including the modifier allele, the modifier spreads rapidly to fixation in asexual strains, because it is associated with genomes that have fewer TE copies and therefore a higher relative fitness. As a consequence, TE activity rates decrease in asexual populations (**Fig. S3**). By contrast, the modifier cannot spread as rapidly in sexual populations because recombination constantly breaks up the association between the modifier and less TE loaded backgrounds. By allowing for the evolution of TE activity rates in our simulations, we were able to identify parameter values representative for yeast that result in simulations with a very close fit to our empirical results (**Fig. 1B, Table S3**). These analyses thus corroborate our empirical findings that a likely mechanism driving genomic TE load reduction in asexual yeast strains is the rapid evolution of increased TE excision rates. A similar effect would be expected if our modifier acted on transposition rather than excision rates, since the net TE activity depends on the relative rates of transposition vs excision. However, our empirical results do not suggest major differences in transposition rates between sexual and asexual yeast strains. In combination with our findings that in the absence of TE activity evolution, sexual strains always lose TEs faster than asexual ones, the empirical results are best explained by an increase in TE excision rates under asexuality (**Fig. 1, 2**).

Our study conclusively shows that sexual reproduction drives the maintenance of TE load in *S. cerevisiae*, while in its absence, TE loads decrease, likely via the evolution of TE activity rates. These findings are consistent with the idea that TEs should evolve to be benign in asexual species, because the evolutionary interests of TEs and their host genome are aligned (Charlesworth and Langley, 1986). While the exact mechanisms causing TE activity change in the asexual yeast populations cannot be assessed in the empirical data, our simulations suggest that there is some form of TE defense mechanism (a ‘modifier locus’) that either segregates in the ancestral yeast strain used in the experiments or repeatedly appeared *de novo* during experimental evolution. Independently of the exact mechanism underlying TE activity evolution in asexual populations, we show that TE loads do not increase, but decrease, in asexual populations. This contrasts with the hypothesis that most asexual species are evolutionarily short lived because they are driven to extinction via negative consequences of accumulating TE copies (Arkhipova and Meselson, 2005; Nuzhdin and Petrov, 2003). Instead, sex, which is the main form of reproduction in eukaryotes, is at the root of the evolutionary success of parasitic TEs.

## Methods

### Yeast experimental evolution

We used data generated in a previous study based on experimental evolution of the yeast *S. cerevisiae* (for in-depth details see (McDonald et al., 2016). In short, 12 different strains were initiated from the same pool of ancestral strains (derived from haploid W303 strains) and kept under constant conditions. Sexual reproduction in yeast depends on the presence of two separate mating types. Only individuals with different mating types can fuse and go through meiosis. Asexual reproduction occurs through budding. For the experiment, six strains consisting of mating type a (MATa) and six of mating type α (MATα), were grown over 990 generations. Of these, four strains were grown exclusively asexually (two of MATa, two of MATα), while the eight others (four of MATa, four of MATα) were mixed for mating events every 90 generations, resulting in four sexual strains. Paired-end Illumina reads were generated for each of the 12 different strains every 90 generations during 990 generations (for a total of 11 sequencing events per strain). Read numbers per sample ranged from 12,775 to 10,270,312, averaging 2,964,869 reads per sample, with a total of 818,303,966 reads. Details of the read data can be found at BioProject PRJNA308843 and in the original study (McDonald et al., 2016).

### Data processing

The genome of the haploid W303 *S. cerevisiae* strain was retrieved from (Lang et al., 2013). All Illumina paired-end raw reads of the 12 replicate strains generated in (McDonald et al., 2016) were downloaded from the SRA (BioProject identifier PRJNA308843). Raw reads were quality filtered by first removing adapter sequences (with the script used in the original study (McDonald et al., 2016), provided by Daniel P Rice, Harvard University), followed by removing the first 10 bases and quality trimming using trimmomatic v0.33 (Bolger et al., 2014) with parameters set to LEADING:3 TRAILING:3 HEADCROP:10 SLIDINGWINDOW:4:15 MINLEN:36. Additionally, non-overlapping paired-reads were constructed *in silico* from the subset of the original paired-reads that were overlapping, as a prerequisite to run the insertion detection pipeline. For this, overlapping reads (on average overlapping by 16 bp) were merged using PEAR v0.9.6 with standard parameters (Zhang et al., 2014). Merged reads were split in half and 20 bp deleted from each read at the overlapping ends using the fastx_toolkit v 0.0.13.2 (Hannon Laboratory, 2010). This resulted in mean read lengths of 72 bp. These ‘artificial’ non-overlapping read pairs were afterwards merged with the read set fraction that was non-overlapping.

### Overall transposable element load

A *S. cerevisiae* specific, curated and updated TE library that contained all consensus sequences of all TE families found in this species is available from (Carr et al., 2012). With this library, we identified TE content and specific copy insertions in the W303 genome using RepeatMasker v4.02 (Smit et al., 2013-2015) with parameters set to -nolow -gccalc -s -cutoff 200 -no_is -nolow -norna -gff -u -engine rmblast. For overall TE load estimates, the fraction of reads mapped to TEs out of total mappable reads was calculated. For that, the TE library was appended to the masked W303 genome and all reads for all strains and generations were mapped using BWA v0.7.13 with standard parameters (Li, 2013). For all strains, mean per-base coverage was checked with bedtools genomecov v2.26 (Quinlan and Hall, 2010), upon which the asexual strain 3D-90 was excluded from all further analyses, as coverage was lower than one-fold for this sample. Following, stat-reads from the PopoolationTE2 v1.10.04 program (Kofler et al., 2016) was utilized to extract the number of total mapped reads and reads mapped to TEs.

### Specific transposable element insertions

To detect specific reference (present in the reference genome) and non-reference TE insertions in all samples, the McClintock pipeline was utilized (Nelson et al., 2017). This pipeline combines six different, benchmarked programs in a standardized fashion. McClintock was run with the non-overlapping read set, the curated TE library, and the W303 assembly using default parameters. The nonredundant insertions output file per sub-program was collected. Next, we utilized a custom python script to collect all information on insertions detected by all different programs and counted insertions with evidence from different programs only once.

To identify full-length TEs and solo LTR insertions, we tagged insertions by length according to the typical *TY* TE properties found in *S. cerevisiae* (i.e. a full TE is a combination of internal sequence and two LTRs within a 500 bp range; solo LTRs are between 220 and 420 bp; see **table S1**). Because TE insertion detection was influenced by coverage, it had to be taken into account for calculating the number of insertions, by adding coverage as random factor (coverage effect *P* < 0.001, generation effect *P* = 0.006, reproductive mode effect *P* = 0.033, and interaction between generation and mode *P* < 0.001; permutation ANOVA). We then calculated the number of lost TEs in asexual strains from the regression slope in asexuals after correcting for coverage (residuals) over 1000 generations, assuming 50 full-length TEs in the ancestor. To additionally check for a bias due to coverage differences between sexual and asexual strains, we randomly subsampled read data for each sample corresponding to the mean coverage of the asexual strains for each generation.

### Modelling

To model TE dynamics in yeast we adjusted an individual based, forward in time simulator by Dolgin and Charlesworth (Dolgin and Charlesworth, 2006). We extended the model to include sexual cycles via fusion of two haploid individuals and recombination, with on average one cross-over on each of the 16 modeled chromosomes (yeast has 16 chromosomes; Goffeau et al., 1996; McDonald et al., 2016). Each chromosome carries 200 loci that are potential targets for a TE insertion. A simulation is initiated with a single individual with 50 TEs randomly placed in the 3200 loci of the genome. The founder individual then populates clonally the whole simulated deme of 100,000 individuals. To account for mutations during this phase we ran 20 burn-in generations of transposition and excision cycles on every individual separately without applying selection. One generation in the simulation consists of a round of selection and reproduction where transposition happens during reproduction, followed by excision. The relative fitness *w*_*n*_ of an individual carrying *n* TEs was modeled as 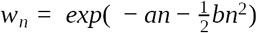, where *a* and *b* are parameters modeling the strength of selection and the strength of epistatic interactions between TEs respectively (Dolgin and Charlesworth, 2006). The simulation was then continued for 990 generations. We performed 10 replicates of each simulation. Using average TE load in the population calculated every ten generations, we fit a linear model to estimate average TE loss across the ten replicates of each simulation. Parameters were derived from yeast experimental measurements and simulations were run with perturbation in the surrounding parametric space (see **table S2**). We further explored the effects of different transposition rates during meiosis vs asexual reproduction, but this did not change the dynamics even for meiotic transposition rates that were not biologically relevant (up to 0.1, i.e. 10% of TEs have transposed during meiosis). The last extension included the introduction of an unliked, general modifier allele increasing the excision rates. The parameters related to this extension are the initial frequency of the modifier allele and the excision rate increases when the modifier allele is present (see **table S3**). See the code documentation for details.

## Acknowledgments

We thank Michael J McDonald, Daniel P Rice and Michael M Desai for providing the experimental evolution raw data and for helpful explanations. We further thank Patrick Tran Van for setting up the insertion pipeline, Daniel L Jeffries for providing the TE wrapper script, Beatriz Navarro Dominguez for improving the empirical analyses R script and Laurent Keller for discussions and comments on the manuscript. This study was supported by a DFG research fellowships (grant numbers BA 5800/1-1 and BA 5800/2-1 to J.B.) and by funding from the University of Lausanne and Swiss SNF (grant numbers PP00P3_170627 and PP00P3_139013 to T.S.).

## Author contributions

J.B. and T.S. conceived the study, J.B. and D.S. performed empirical data analyses, K.S.J. and D.R performed modeling, J.B. and T.S. wrote the paper with input from all authors.

## Data availability

Raw read data of the experiment is available at SRA (BioProject identifier PRJNA308843).

## Code availability

The code used for both the analyses of empirical data and for the theoretical prediction of TE dynamics together with explanations are available online at https://github.com/KamilSJaron/reproductive_mode_TE_dynamics

## Supplementary information

**Fig. S1.**
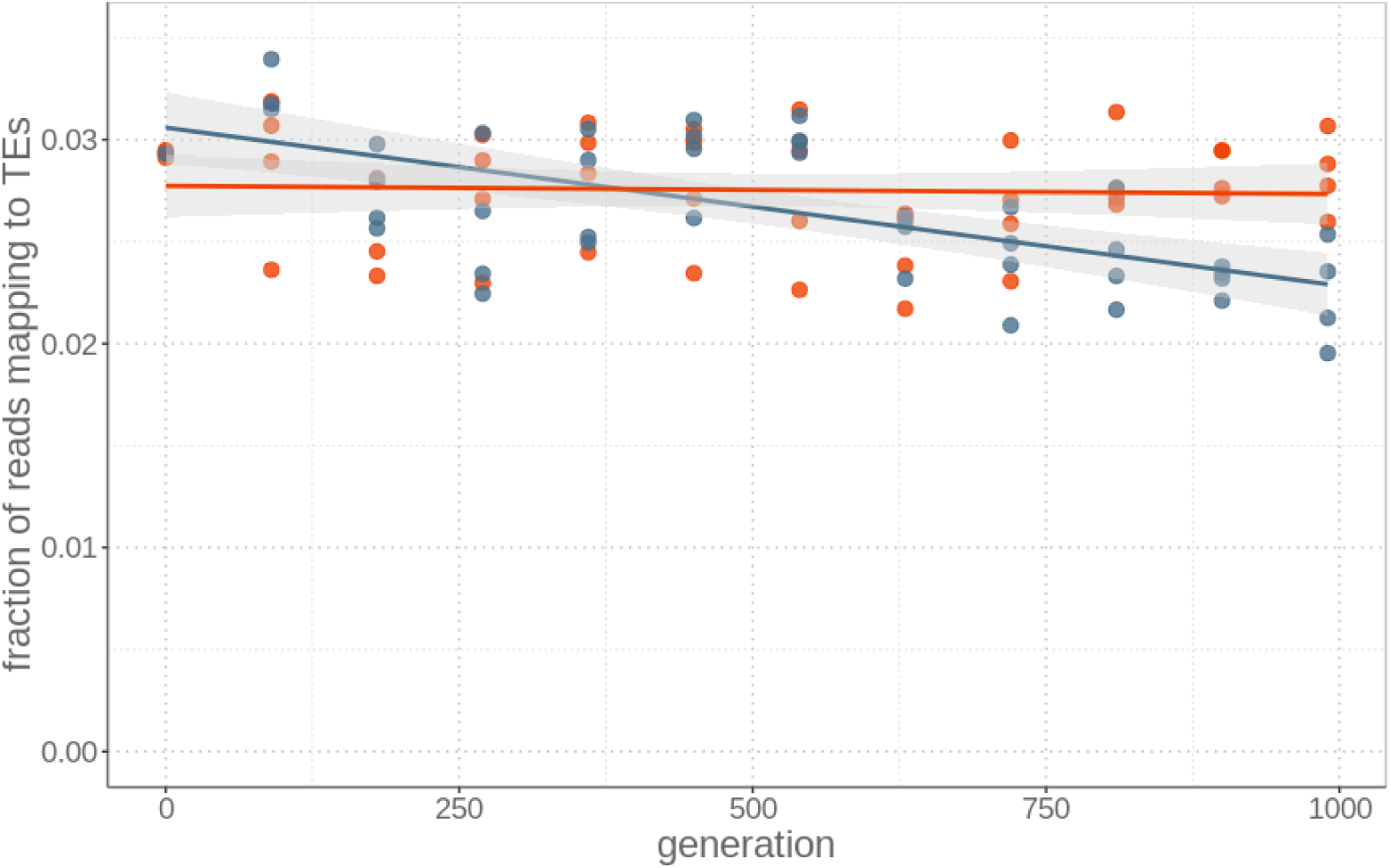
Transposable element load remains stable in sexual strains, but is reduced in asexual strains after 1000 generations. Read fraction mapping to TEs relative to the sum of reads mapping to the genome and/or the TE library for each of the four replicate sexual (red) and asexual (blue) strains sequenced every 90 generations (from generation 0 to 990). Lines represent LR and the grey areas represent 95% CI.

**Fig. S2.**
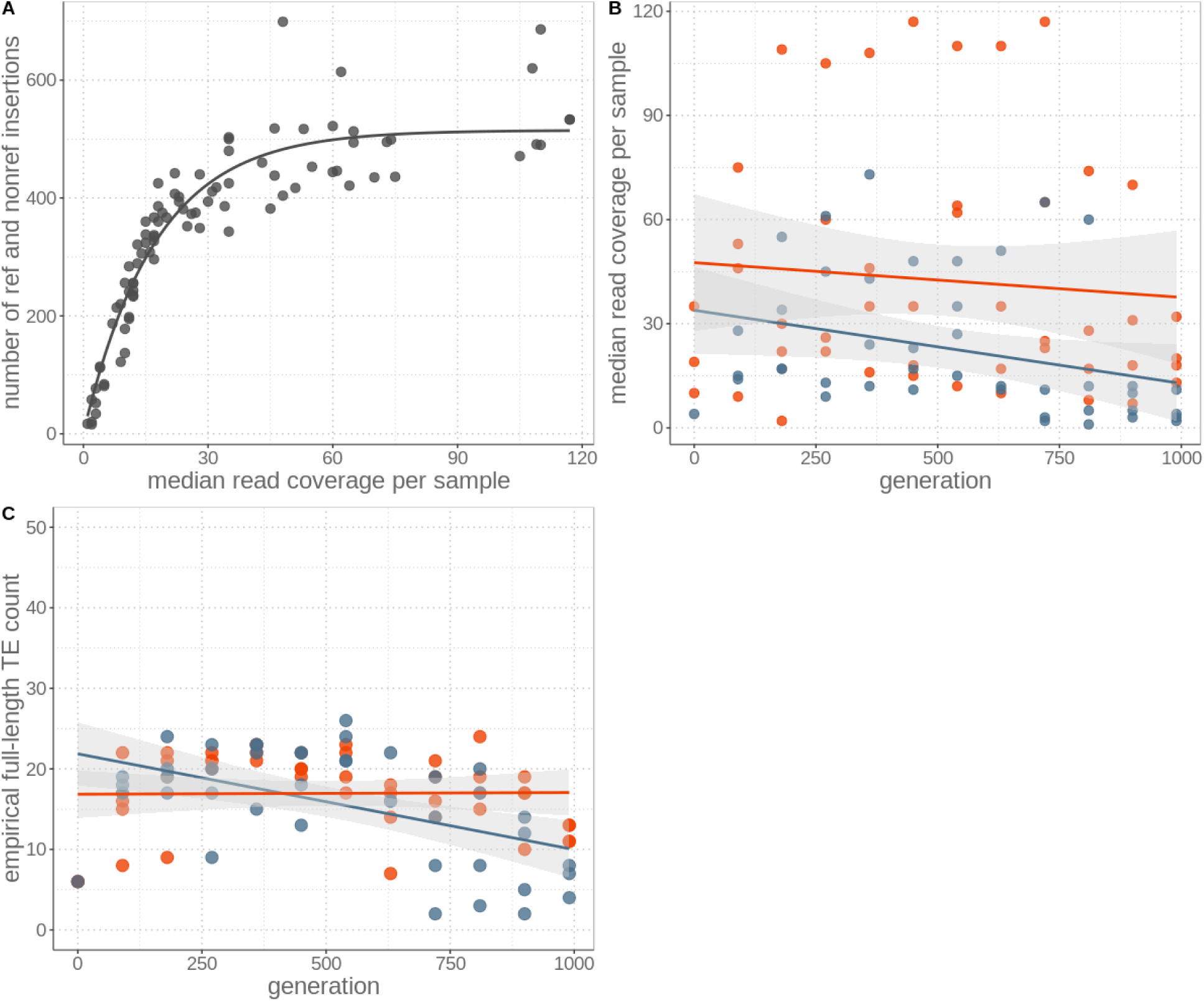
Identification of TE insertions depends on the sequencing coverage. (A) TE insertions (including those present in the reference genome and *de novo* insertions) *vs*. median sequencing coverage from paired reads. Coverage influences the ability to detect TE insertions (Wilcoxon signed-rank test V = 4095, p-value < 0.001). (B) Median read coverage per sample for sexual (red) and asexual (blue) strains over 1000 generations. Data from asexual strains had lower coverage, but were not different to sexuals through time (generation effect *P* = 0.096, reproductive mode effect *P* = 0.002, and interaction between generation and mode *P* = 0.588; permutation ANOVA). Lines represent LR and the grey areas represent 95% CI. (C) Subsampling to the mean asexual read coverage per generation for all samples results in similar findings (generation effect P = 0.012, reproductive mode effect P = 0.302, and interaction between generation and mode P = 0.004; permutation ANOVA).

**Fig. S3.**
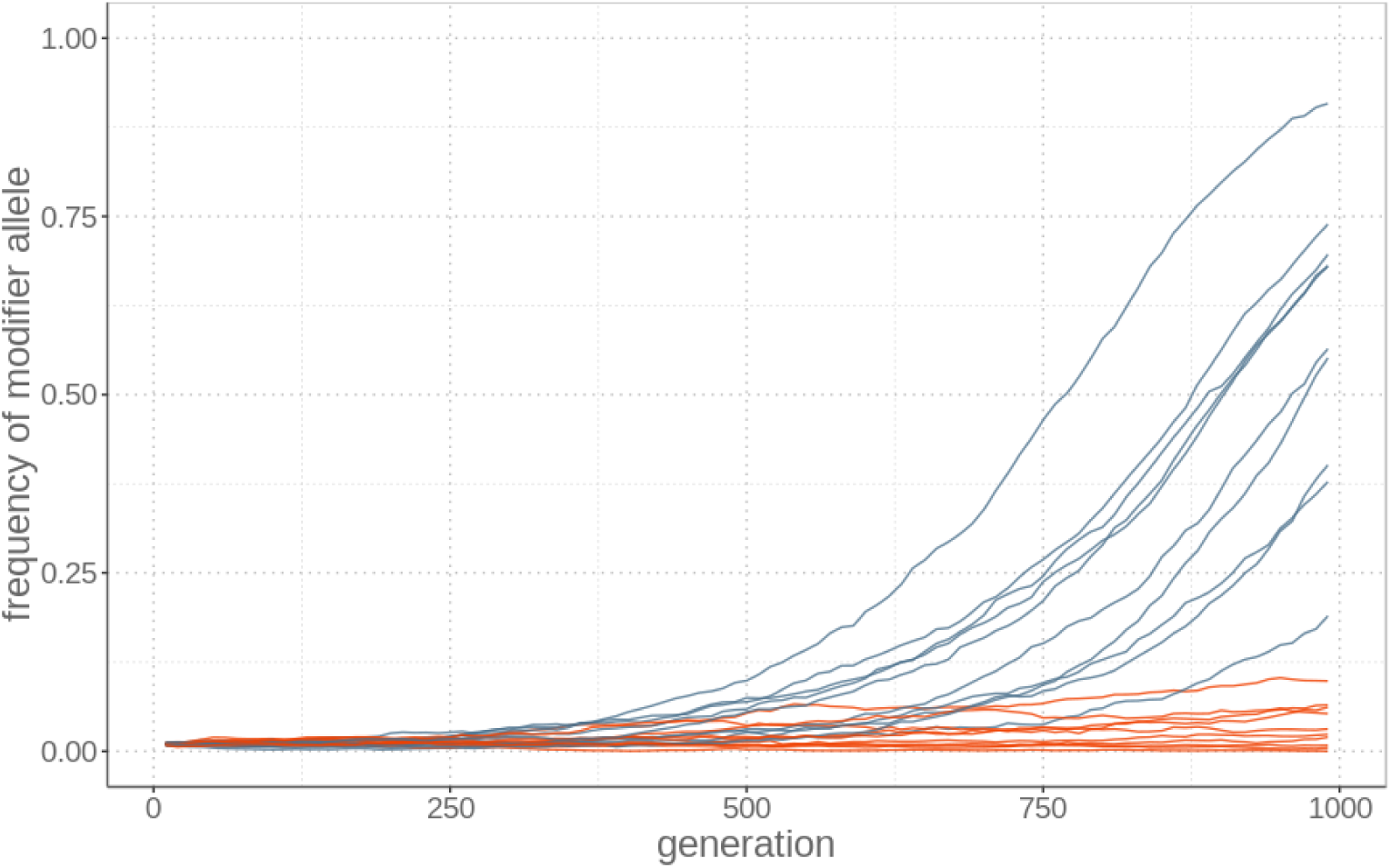
The spread of a modifier of excision rates is faster in asexual than sexual populations because it remains linked to genomes that have few TE copies and therefore a high relative fitness. The modifier allele frequency is shown over time for simulations under sexual (red) and asexual (blue) reproduction, with ten replicates.

**Fig. S4.**
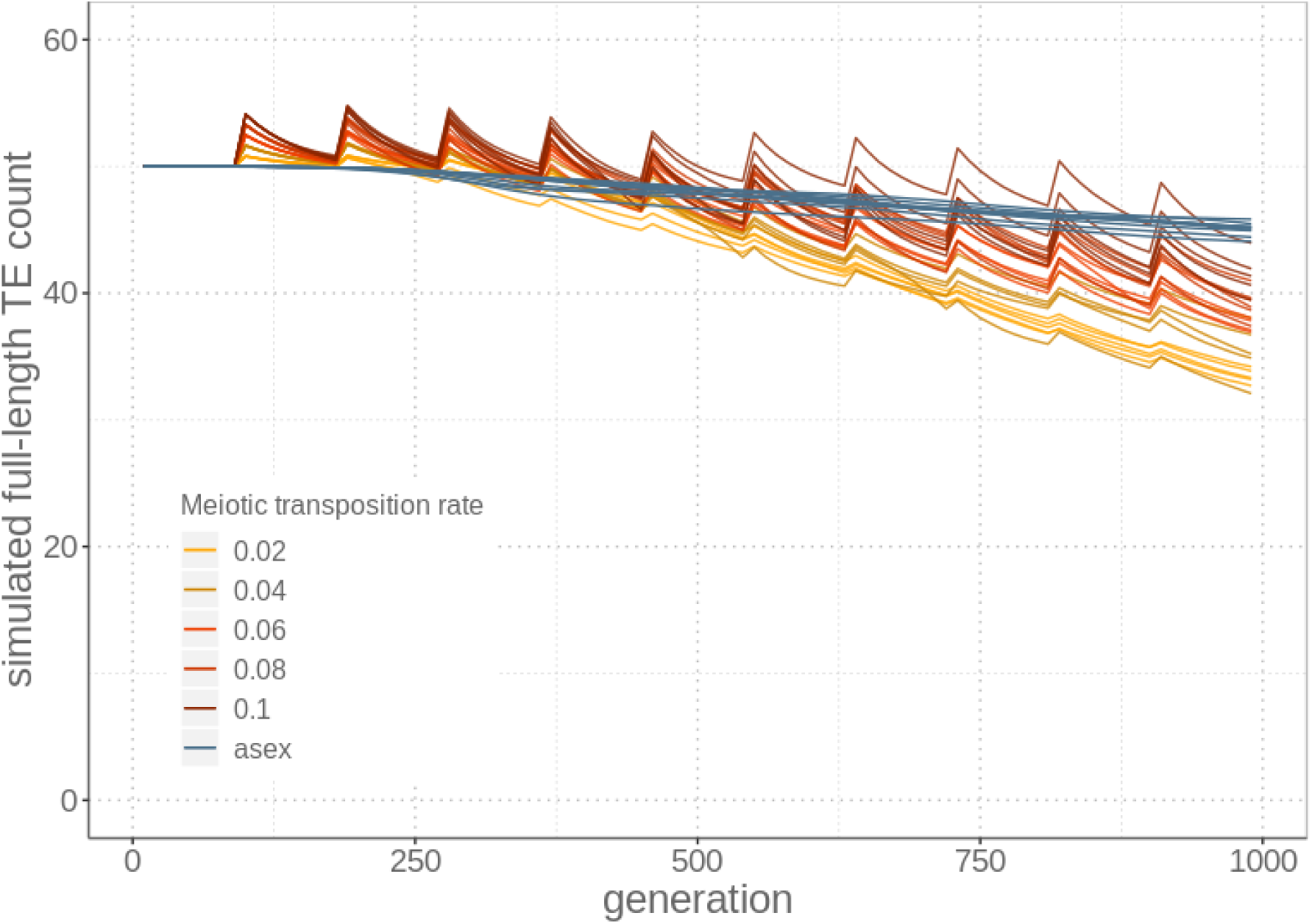
Simulations with higher transposition rates during meiosis than mitosis. Meiosis generates the TE load spikes following events of sexual reproduction, but allows for selection to effectively remove genotypes with high TE loads by generating fitness variation among genotypes. Parameters used in the simulations are indicated in table S2 (bold values).

**Table S1.**
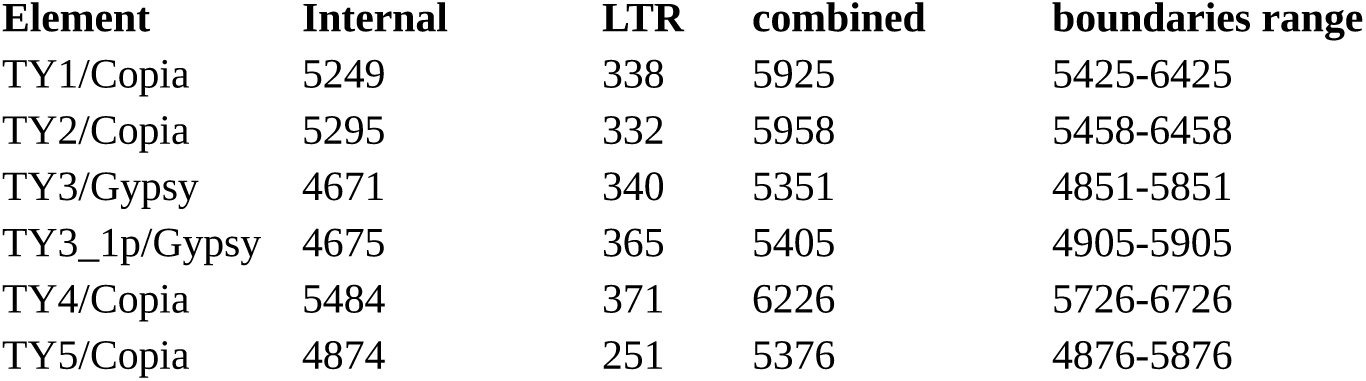
*S. cerevisiae* TY elements and the sizes (in bp) of internal regions and LTRs and the size boundaries used for filtering.

**Table S2.** Explored parameter space of the simulations as pertinent for yeast (empirically determined values in bold). Selection_a and selection_b are selection coefficients for linear fitness effects and epistasis, respectively. Lost_TEs refers to the total number of TE lost after 1000 generations (averaged over ten replicates).

| transposition_rate | exision_rate | selection_a | selection_b | sex_lost_TEs | asex_lost_TEs |
| --- | --- | --- | --- | --- | --- |
| 1.00E-06 | 5.00E-07 | 1.00E-04 | 0 | 6.5 | 2 |
| 1.00E-05 | 5.00E-07 | 1.00E-04 | 0 | 6.7 | 1.5 |
| 1.00E-04 | 5.00E-07 | 1.00E-04 | 0 | 3.5 | -0.8 |
| 1.00E-06 | 1.00E-06 | 1.00E-04 | 0 | 8.1 | 2.2 |
| 1.00E-05 | 1.00E-06 | 1.00E-04 | 0 | 7.8 | 2.3 |
| 1.00E-04 | 1.00E-06 | 1.00E-04 | 0 | 5.7 | -0.1 |
| 1.00E-06 | 5.00E-05 | 1.00E-04 | 0 | 28.6 | 17 |
| 1.00E-05 | 5.00E-05 | 1.00E-04 | 0 | 21.9 | 13.1 |
| 1.00E-04 | 5.00E-05 | 1.00E-04 | 0 | 19.4 | 11 |
| 1.00E-06 | 5.00E-07 | 5.10E-04 | 0 | 6.7 | 1.9 |
| 1.00E-05 | 5.00E-07 | 5.10E-04 | 0 | 6.4 | 1.6 |
| 1.00E-04 | 5.00E-07 | 5.10E-04 | 0 | 4.3 | -0.2 |
| 1.00E-06 | 1.00E-06 | 5.10E-04 | 0 | 9.1 | 2.5 |
| 1.00E-05 | 1.00E-06 | 5.10E-04 | 0 | 8.2 | 2.4 |
| 1.00E-04 | 1.00E-06 | 5.10E-04 | 0 | 6.3 | 0.5 |
| 1.00E-06 | 5.00E-05 | 5.10E-04 | 0 | 22.7 | 14 |
| 1.00E-05 | 5.00E-05 | 5.10E-04 | 0 | 22.6 | 13.6 |
| 1.00E-04 | 5.00E-05 | 5.10E-04 | 0 | 20.6 | 11.3 |
| 1.00E-06 | 5.00E-07 | 1.00E-04 | 0.00039 | 6.5 | 2 |
| 1.00E-05 | 5.00E-07 | 1.00E-04 | 0.00039 | 6.7 | 1.5 |
| 1.00E-04 | 5.00E-07 | 1.00E-04 | 0.00039 | 3.5 | -0.8 |
| 1.00E-06 | 1.00E-06 | 1.00E-04 | 0.00039 | 8.1 | 2.2 |
| 1.00E-05 | 1.00E-06 | 1.00E-04 | 0.00039 | 7.8 | 2.3 |
| 1.00E-04 | 1.00E-06 | 1.00E-04 | 0.00039 | 5.7 | -0.1 |
| 1.00E-06 | 5.00E-05 | 1.00E-04 | 0.00039 | 28.6 | 17 |
| 1.00E-05 | 5.00E-05 | 1.00E-04 | 0.00039 | 21.9 | 13.1 |
| 1.00E-04 | 5.00E-05 | 1.00E-04 | 0.00039 | 19.4 | 11 |
| 1.00E-06 | 5.00E-07 | 5.10E-04 | 0.00039 | 6.7 | 1.9 |
| 1.00E-05 | 5.00E-07 | 5.10E-04 | 0.00039 | 6.4 | 1.6 |
| 1.00E-04 | 5.00E-07 | 5.10E-04 | 0.00039 | 4.3 | -0.2 |
| **1.00E-06** | **1.00E-06** | **5.10E-04** | **0.00039** | **9.1** | **2.5** |
| 1.00E-05 | 1.00E-06 | 5.10E-04 | 0.00039 | 8.2 | 2.4 |
| 1.00E-04 | 1.00E-06 | 5.10E-04 | 0.00039 | 6.3 | 0.5 |
| 1.00E-06 | 5.00E-05 | 5.10E-04 | 0.00039 | 22.7 | 14 |
| 1.00E-05 | 5.00E-05 | 5.10E-04 | 0.00039 | 22.6 | 13.6 |
| 1.00E-04 | 5.00E-05 | 5.10E-04 | 0.00039 | 20.6 | 11.3 |

**Table S3.** Explored parameter space for simulations including a modifier allele. Highlighted is the simulation closest to empirical observations. Init_f is the frequency of the modifier at the start of the simulations. Selection_a and selection_b are selection coefficients for linear fitness effects and epistasis, respectively. Lost_TEs refers to the total number of TE lost after 1000 generations (averaged over ten replicates). The bold lines refer to parameter combinations that generate results close to the observed empirical values.

| init_f | selection_a | selection_b | sex_lost_TEs | asex_lost_TEs |
| --- | --- | --- | --- | --- |
| 0.01 | 2.00E-04 | 0 | 0.6 | 1.2 |
| 0.01 | 3.00E-04 | 0 | 0.7 | 2.7 |
| 0.01 | 4.00E-04 | 0 | 0.6 | 5.3 |
| **0.01** | **0.000425** | **0** | **0.9** | **6.2** |
| **0.01** | **0.00045** | **0** | **0.7** | **6.6** |
| **0.01** | **0.000475** | **0** | **0.8** | **10.3** |
| **0.01** | **5.00E-04** | **0** | **1** | **9.9** |
| 0.01 | 5.00E-04 | 1.00E-06 | 0.3 | 0.3 |
| 0.01 | 1.00E-03 | 1.00E-06 | 0.6 | 0.9 |
| 0.01 | 2.00E-03 | 1.00E-06 | 1.3 | 5.1 |
| 0.01 | 2.00E-04 | 1.00E-05 | 1.3 | 15.6 |
| 0.01 | 3.00E-04 | 1.00E-05 | 1.2 | 17 |
| 0.01 | 4.00E-04 | 1.00E-05 | 2.2 | 20.1 |
| 0.01 | 5.00E-04 | 1.00E-05 | 2.2 | 22.7 |
| 0.1 | 2.00E-04 | 0.00E+00 | 3.4 | 7.6 |
| 0.1 | 3.00E-04 | 0.00E+00 | 4 | 12.6 |
| 0.1 | 4.00E-04 | 0.00E+00 | 4.9 | 16.9 |
| 0.1 | 5.00E-04 | 0 | 6.4 | 20 |
| 0.1 | 2.00E-04 | 1.00E-05 | 7.6 | 22.4 |
| 0.1 | 3.00E-04 | 1.00E-05 | 8.8 | 23.5 |
| 0.1 | 4.00E-04 | 1.00E-05 | 10.6 | 24.9 |
| 0.1 | 5.00E-04 | 1.00E-05 | 11.9 | 26.1 |

